# Targeted mutagenesis of the SUMO protease, *Overly Tolerant to Salt1* in rice through CRISPR/Cas9-mediated genome editing reveals a major role of this SUMO protease in salt tolerance

**DOI:** 10.1101/555706

**Authors:** Cunjin Zhang, Anjil Kumar Srivastava, Ari Sadanandom

## Abstract

SUMO proteases are encoded by a large gene family in rice and are a potential source of specificity within the SUMO system that is responding to different environmental cues. We previously demonstrated a critical role of OsOTS class of SUMO proteases in salt and drought stress in rice by silencing several family members collectively via RNAi methods. However, to date it has not been possible to assign a role to specific family members due to limitations of RNAi mediated off target silencing across several members of the gene family. The clustered regularly interspaced short palindromic repeats (CRISPR)-associated endonuclease 9 (CRISPR/Cas9) system has emerged as a promising technology for specific gene editing in crop plants. Here, we demonstrate targeted mutagenesis of *OsOTS1* in rice using the CRISPR/Cas9 gene editing system in the rice cultivar Kitaake. Guide RNA mediated mutations in *OsOTS1* was highly efficient with almost 95% of T_0_ transgenics showing the desired effect with no off-target mutations. The *OsOTS1* mutations observed in T_0_ generation were heritable in subsequent generations. *OsOTS1* CRISPR lines show enhanced sensitivity to salt with reduced root and shoot biomass indicating that OsOTS1 has a major role in salt stress tolerance in rice. This unexpected finding indicates that precise and effective genome editing can be used to characterise specificity within the SUMO system in rice.

---

Genome editing has opened up new avenues to engineer desirable biological systems. Due to its simplicity, high efficiency and design flexibility, the CRISPR/Cas9 (clustered regularly interspaced short palindromic repeats–associated nuclease 9) system is widely used for genome editing in various plants and can be employed to discover non-redundant functions in plant gene families^1–2^. We wanted to use CRISPR gene editing to understand the role of specific components of SUMOylation, an emerging post-translational modification (PTM) system in plants.

SUMO is covalently attached to protein targets through an isopeptide bond that occurs through a lysine amino acid on the protein target and the SUMO C-terminus, in a process analogous to that of ubiquitin activation and conjugation. Prior to activation SUMO is proteolytically processed from its precursor form to produce a conserved C-terminal Gly-Gly motif. SUMO is catalysed by different sets of enzymes to facilitate direct transfer of SUMO to a lysine amino group within the protein target. The accumulation of SUMO conjugated proteins in the cell is in part counterbalanced by dedicated SUMO-specific cysteine proteases (SUMO proteases) that cleave SUMO off their targets. SUMO deconjugation is equally important and crucial to the balance between SUMOylated and non-SUMOylated proteins. The analysis of different SUMO proteases is beginning to reveal complex patterns of substrate specificity amongst the SUMO proteases in model plants and crops^3^.

Rice is the major staple cereal crop across the world and consumed by half of the world population. Tolerance to salt stress is a major factor in rice productivity^4^. Salinity interferes with growth and development of rice plants. We previously characterized and demonstrated the critical role of OsOTS class of SUMO proteases in salt and drought stress in rice^5–6^. We demonstrated that high levels of SUMO conjugated proteins accumulate in *OsOTS*-RNAi rice plants during salt stress lead to growth retardation. However, in non-salt conditions they are morphologically indistinguishable from wild type plants. Transgenic rice plants overexpressing *OsOTS1* overexpressing plants are more salt tolerant with a concomitant reduction in the levels of SUMOylated proteins. We have also found that overexpression of *OsOTS1* confers salt tolerance in rice by increasing root biomass. In high salt stress conditions, the degradation rate of OsOTS1 protein is enhanced revealing that SUMO conjugation in rice plants during salt stress is in part achieved by down-regulation of OTS1/2 activity^5^. OsOTS1 SUMO protease also directly targets the ABA and drought responsive transcription factor OsbZIP23 for deSUMOylation affecting its stability. OsOTS-RNAi lines show increased abundance of OsbZIP23 and increased drought responsive gene expression while OsOTS1-OX lines show reduced levels of OsbZIP23 leading to the suppression of drought responsive gene expression^6^. However, the development of gene function research is restricted due to the off-target effect and the incompleteness of regulation via RNA interference. Therefore, new gene knock-out technologies with higher specificity and lower off target rate are necessary to uncover non-redundant functions in gene families.

In this study, we applied the CRISPR/Cas9 gene editing system to produce knockouts of the SUMO protease, *OsOTS1* gene in the rice cultivar Kitaake, which will provide an excellent foundation to analyse the functional significance of this specific SUMO protease in amongst the OsOTS SUMO protease gene family. Introduction of guide RNA mediated mutations in *OsOTS1* was highly efficient with almost 95% of T_0_ transgenics showing the desired mutation with no off-target mutations. The OsOTS1 mutations observed in T_0_ generation were heritable in subsequent generations. *OsOTS1* CRISPR lines show enhanced sensitivity to salt with reduced root and shoot biomass indicating that OsOTS1 alone has a major role in salt stress tolerance in rice. This unexpected finding indicates that precise and effective genome editing can be used to characterise specific SUMO protease gene function in rice.

## Results

### CRISPR/Cas9-mediated mutation of OsOTS1 in rice

To exploit the CRISPR/Cas9 system to induce mutations at the *OsOTS1* gene, we customised a sgRNA to target the SUMO protease gene *OsOTS1* in rice (Fig. 1). To test whether the customised sgRNA specifically targets rice OTS1, we first verified the *Cas9* cassette in transgenic Kitaake rice cultivar by PCR to detect the gene encoding Hygromycin B phosphotransferase that confers resistance to the antibiotic hygromycin; The presence of HYG gene have been confirmed for all 15 transgenic lines by PCR. Then we identified the target gene mutation in these transgenic lines by DNA sequence analysis (Fig. 2a). In total, 14 individual rice transgenic T_0_ lines were subjected to mutation detection using Sanger sequencing of PCR products amplified with primers flanking the sgRNA target sites from leaf tissue. The total of three different kinds of mutations were detected. The genomic DNA analysis from T_0_ seedling leaves revealed the following different mutations: 1-bp insertion, 1 or 2-bp deletion and 5-bp deletions (Fig. 2b). In each case either insertion or deletions in *OsOTS1* gene is predicted to lead to frameshifts and premature stops producing aberrant proteins.

**Figure 1.**
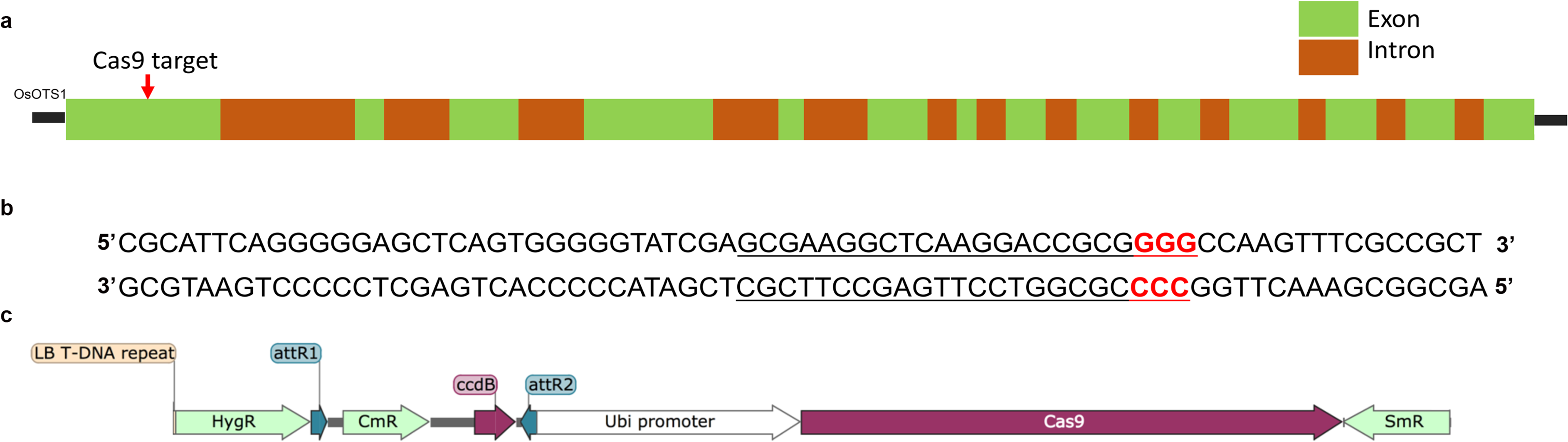
Schematic of the design of CRISPR guide RNA for targeting *OsOTS1* gene in rice. **a** Gene structure of the *OsOTS1* gene. The target site of Cas9 located in the coding sequence of the first exon. b SgRNA target site for *OsOTS1.* Red font indicates PAM sequence. c Schematic illustration of CRISPR/ Cas9 DNA construct used in this study. HygR = Hygromycin resistance CmR = Chloramphenicol resistance, SmR = Spectinomycin resistance, Ubi Promoter = Ubiquitin promoter.

**Figure 2.**
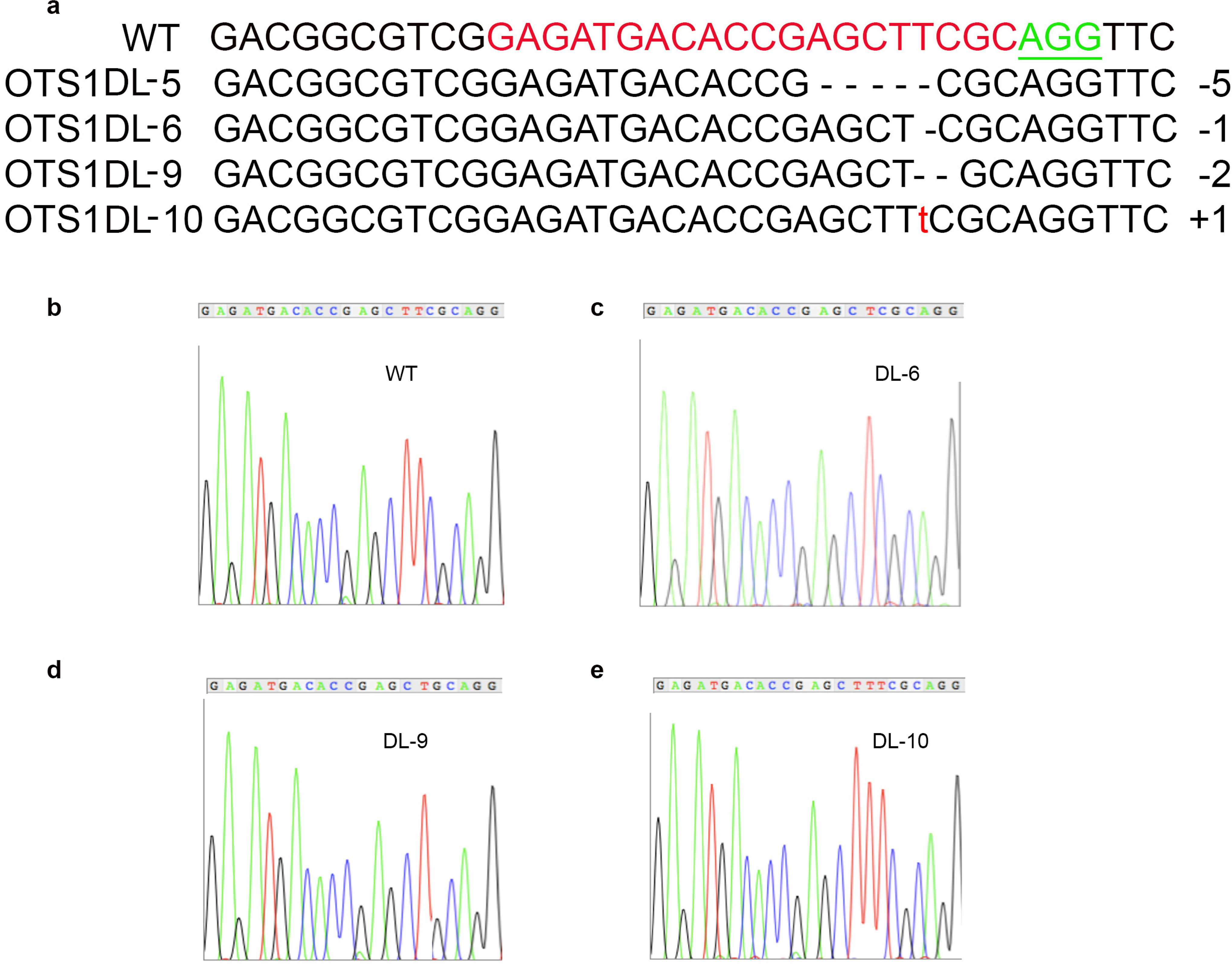
DNA Sequence analysis reveals targeted mutagenesis of OsOTS1 in rice using the CRISPR/Cas9 system. a DNA sequence of wild type and selected transgenic lines indicating deletions and insertion in the *OsOTS1* gene. Red bases indicate targeting sequences for sgRNA; Green bases indicate PAM sites. b-e Sanger Sequencing chromatograms of DNA from wild type and the homozygous mutant Os0TS1 CRISPR mutants.

### Potential off-targets analysis

Since the rice ULP-like SUMO protease gene family includes 12 members, we assessed the potential off-targets at the other two closely related rice SUMO protease genes, *OsOTS2 and 3*. The genomic regions containing the potential off target site were amplified from wild type and two different mutant lines (DL-6 and DL-10) and the PCR products were sequenced. However, no overlapped indels were detected in these lines. These results indicated that off-targeting did not occur in the candidate sites indicating that DL-6 and DL-10 represent bona fide mutants specifically knocking out OsOTS1.

### Salt sensitive phenotype of OsOTS1 CRISPR rice mutants

In our previous study we used RNAi methods to silence the OTS sub clade of the ULP-like SUMO proteases in rice. This study revealed that silencing this gene sub-clade led to an increase in salt sensitivity in the rice cultivar *Nipponbare*. However, overexpression of *OsOTS1* lead to an increase in salt tolerance in the cultivar *Nipponbare* leading us to speculate that OsOTS1 may have a bigger role in salt tolerance in rice. Therefore, to ascertain this prospect using a loss of function approach we characterised the two different CRISPR mediated OsOTS1 knockout mutant genotypes (DL-6 and DL-10) for salt tolerance in the cultivar kitaake. Seedling growth rates of OsOTS1 SUMO protease knockout lines DL-6 and DL-10 were compared to plants grown on 150mM NaCl as previously described^5^. At the 10-day old seedling stage, no significant growth differences were observed between lines carrying empty vector, *OsOTS-RNAi* and DL-6 and DL-10 *OsOTS1* knockouts under unstressed conditions. However, the same lines grown in ½ MS medium with 150mM NaCl, showed significant growth inhibition indicating severe salt sensitivity (Fig. 3 A-D).

**Figure 3.**
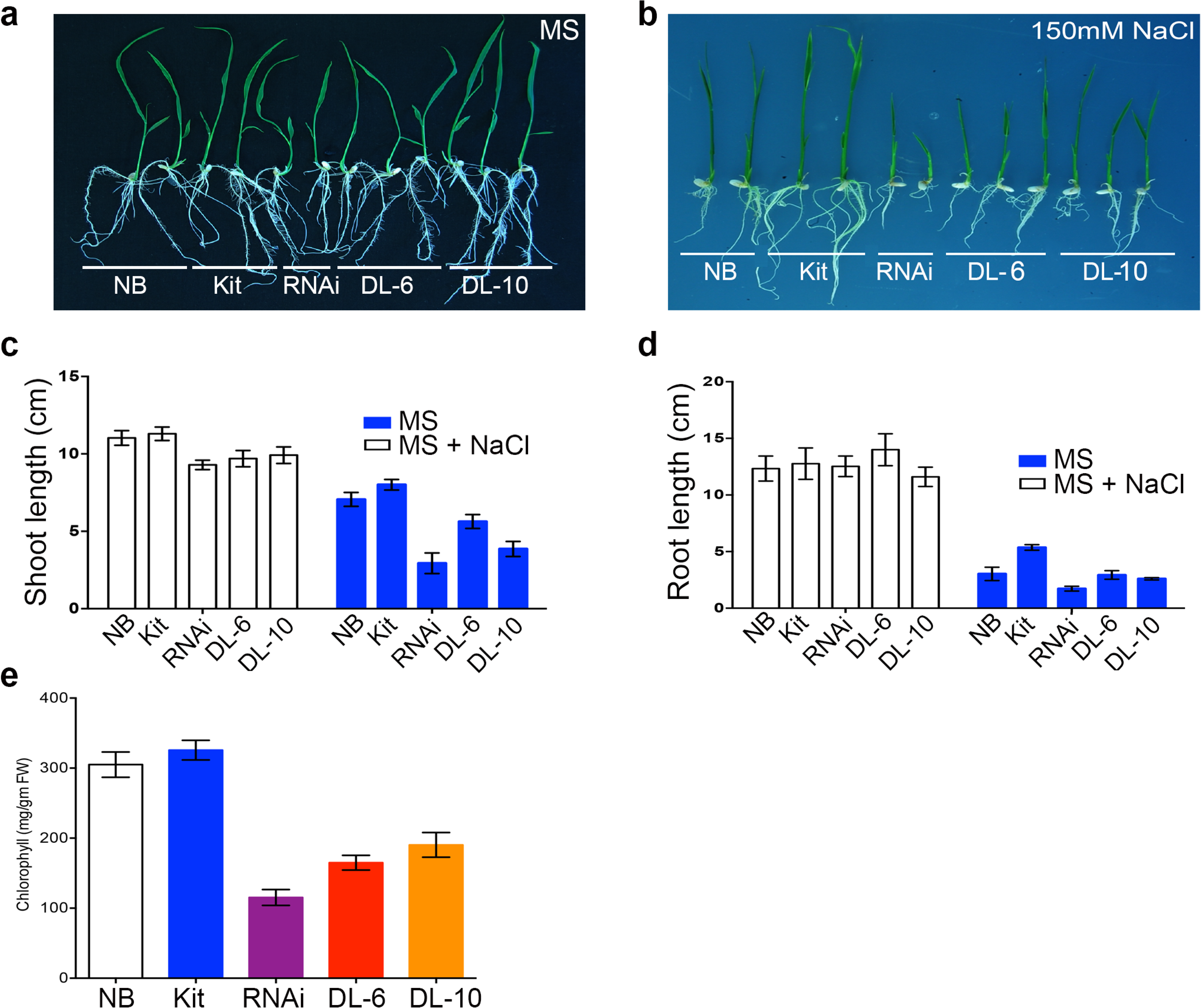
Targeted CRISPR mutation of *OsOTS1* gene in rice results in salt sensitiveity. **a-b** The mutation phenotype of homozygous *OsOTS1* CRSIPR mutant plants. **c-d** Quantification data of shoot and root length for wild type Nipponbare (NB), Kitake (Kit), *Os0TS1* RNAi, CRISPR mutant lines DL-6 and DL-10 line seedlings. **e** Quantification of chlorophyll content at 150mM concentration of salt treatment in various control and homozygous OsOTS1 CRISPR mutants.

One of the key parameters that contribute to reduce growth rate is the inactivation of photosynthesis. This is reflected in the relative chlorophyll content, in plants^7^. Chlorophyll content measurement of seedlings of the various genotypes grown in 150mM salt indicated that DL-6 and DL-10 had greatly reduced chlorophyll content akin to those observed in *OsOTS* RNAi lines when compared to vector only controls (Fig. 3E).

### The expression of OsOTS1 target genes are suppressed in OsOTS1 CRISPR mutants

Quantitative RT-PCR analysis indicated that there was no significant difference in OsOTS1 transcript levels in *DL-6* and *DL-10 OsOTS1* CRISPR mutants compared to wild type Nipponbare and Kitaake (Fig. 4). This is expected as *DL-6* and *DL-10* only contain single base deletions in the *OsOTS1* gene. Although it has been established that OTS sub family of SUMO proteases positively regulates root growth in rice by modulating the gene expression of keys genes that regulate root growth promotion such as *OsDro1* and *OsIAA23*^5^, it remains unknown whether the loss of OsOTS1 alone may have an effect on these genes. To ascertain this, we analysed the transcriptional levels of *OsDro1* and *OsIAA23*^*^8–9^*^ in *OsOTS1* CRISPR mutants along with wild type Nipponbare, Kitaake and RNAi transgenic lines. We found that the transcriptional levels of *OsDro1* and *OsIAA23* was significantly reduced compared to WT controls as well as OsOTS-RNAi lines (Fig. 5). This evidence indicates that OsOTS1 plays an important role in regulating *OsDro1* and *OsIAA23* gene expression therefore affecting root growth. Since DL-6 and DL-10 represent OsOTS1 specific knockouts our data reveals a critical role for OsOTS1 in regulating root growth that cannot be compensated for by OsOTS2 or OsOTS3.

**Figure 4.**
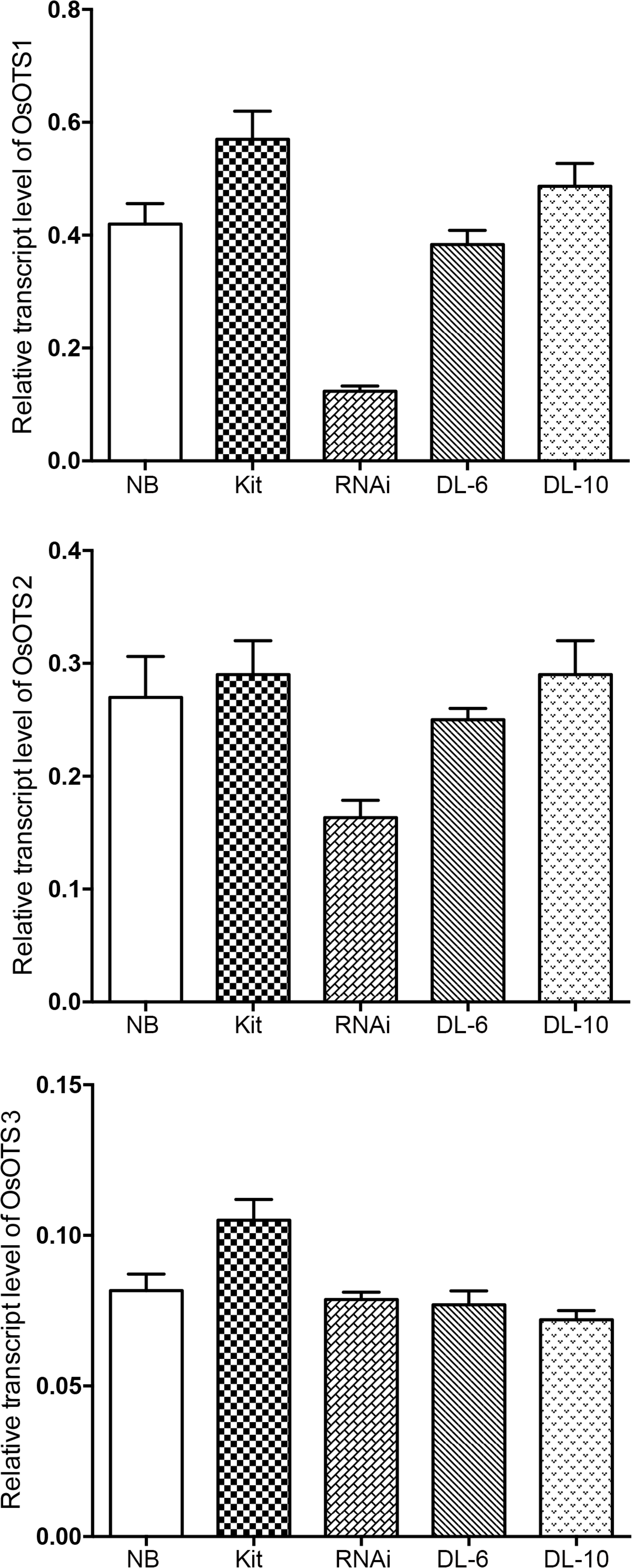
Transcript analysis of *OsOTS1* and its close homologues. **a** q-RT PCR analysis of *OsOTS1* **b** q-RT PCR analysis of *OsOTS2* **c** q-RT PCR analysis of *OsOTS3* in wild type Nipponbare (NB), Kitake (Kit), *OsOTS*-RNAi, and *OSOTS1* CRISPR mutant DL-6 and DL-10 rice seed lings. Total RNA was prepared from the 10-d-old seedlings of the controls and transgenic lines and then reverse transcribed. Transcript levels were measured by qPCR with cDNA. Actin was used as an internal control. Data represent the mean values of three biological replicates. Error bars indicate SD. *P values for difference between control (NB) and the CRISPR mutant: *p<0 .05 (Student’s t-test).

**Figure 5.**
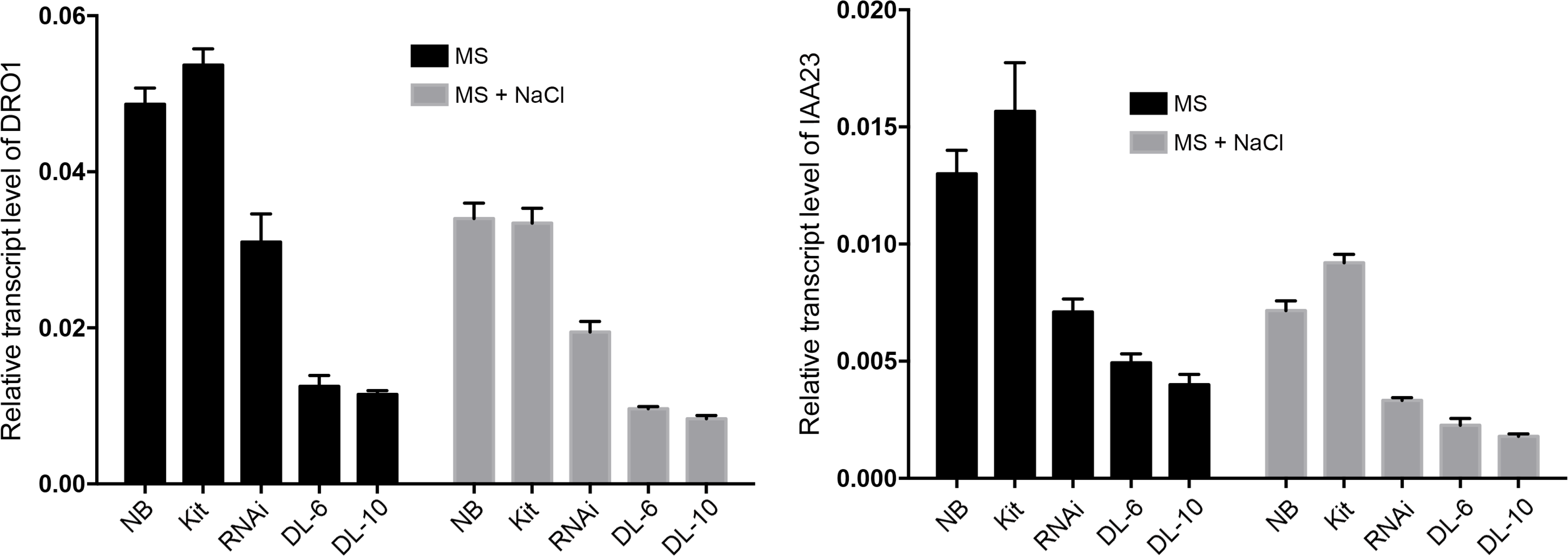
*OsOTS1* affects the expression of root growth promoting genes. Ten-day-old rice seedlings were transferred to MS medium with 150mM NaCl and harvested after 6 h of salt treatment to extract total RNA. The expression of different genes was determined by qPCR. Gene expressions of target genes were normalized to target genes were normalized to that of rice actin. The relative expression was compared with that of control plants. Mean values were obtained from three biologically independent experiments. Error bars indicate SD.

## Discussion

Plant regulatory networks that are important in responding to drought stress overlap considerably with those for other abiotic stresses, including salt stress^10–11^. This evidence indicates that development of drought and salt tolerant crop cultivars is possible and essential for adapting agriculture to climate change^12^. Studies on model plants and crops indicate that yield and quality is critically dependent upon the complex perception and signaling mechanisms that operate to produce an integrated physiological response to environmental stresses^13–14^. Therefore, identifying molecular targets (i.e. that act as ‘Master’ coordinators) that influence multiple stress signaling pathways and revealing how these improve whole plant stress physiology, will be crucial for increasing crop productivity.

PTMs are major regulators of plant stress responses by modulating stress sensors and downstream transcription factors that control the expression of a multitude of stress-related genes. Protein ubiquitination and phosphorylation are the two most understood PTMs controlling stress signaling. In recent years, SUMO (Small Ubiquitin-like Modifier) has emerged as a very influential class of PTMs^15^. A key function of SUMO is to act as a vital counterpoise to ubiquitination, adding a layer of control above ubiquitination with respect to substrate availability, stoichiometry, competition for targets and prevention of ubiquitin dependent protein degradation. Similarly, recent evidence indicates that phosphorylation can be regulated by SUMOylation of kinases and phosphatases indicating a key point of cross talk between the different PTMs^16^. This places SUMOylation as a likely central regulator of signaling in eukaryotes, hence is an ideal target for manipulating complex molecular responses such as salt and drought resistance. Indeed, we have shown that the OsOTS1 SUMO protease gene family is a critical regulator of salt and drought stress in rice using RNAi mediated gene silencing^5^. Previously, we have demonstrated that OsOTS-RNAi rice plants accumulate high levels of SUMO1 conjugated proteins during salt stress and are highly salt sensitive however, in non-salt conditions they are developmentally indistinguishable from wild type plants. Transgenic rice plants overexpressing OsOTS1 have increased salt tolerance and a concomitant reduction in the levels of SUMOylated proteins. However, the impact of individual members of this family has not been established. Here, we demonstrate CRISPR-mediated to generate gene specific knockouts of a closely related SUMO protease without off target effects.

Two independent *OsOTS1* CRISPR knock out mutants did not show any change in expression levels of its target gene. However, based on the salt sensitivity and chlorophyll content analysis, these CRISPR mutants showed increased salt sensitivity compared to empty vector control and this sensitivity was comparable to previously generated RNAi lines. These results suggested that the knockout of *OsOTS1* gene alone is sufficient to generate a significant decrease in the rice salt tolerance. Salt treatment triggers OsOTS1 leading to increased SUMO conjugation in rice plants during salt stress. These results expand our understanding of the role and significance of an individual SUMO protease in salt tolerance in rice. This data suggests that OsOTS2 or 3 has less of an impact on salt stress in rice. However, this does not rule out that OsOTS2 or 3 may have a bigger role in other stresses such as drought. Interestingly we have shown that overexpressing OsOTS1 leads to increased drought sensitivity in pot grown rice plants^7^. This was due to increased root biomass in OsOTS1 rice transgenics that led to greater water loss in pots compared to controls resulting in severe drought during the early growth phase of rice plants. Overexpressing OsOTS1 alone is enough to boost root growth under salinity through upregulation of root growth promoting gene expression via a hitherto unknown mechanism^5^,^17^. In this study *OsOTS1* CRISPR knockouts show a severe downregulation of expression of the root growth promoting genes *OsDRO1* and *OsIAA23*. This down regulation is greater than what was observed in OsOTS1 RNAi lines. The CRISPR loss of function approach further validates the role of OsOTS1 and supports the data generated via the gain of function approach of overexpressing OsOTS1^5^. This evidence demonstrates that OsOTS1 specifically has a major role in regulating the expression of root growth promoting genes in rice. However, the mechanism of how this is achieved is yet to be unravelled.

SUMOylation modifies its target function in many ways, including stimulation of new protein–protein interactions, changing their subcellular localization, stabilizing or marking them for proteasomal degradation^18^. Previously we demonstrated the significance of SUMOylation in mediating OsbZIP23 transcription factor accumulation in rice. This in turn leads to the up-regulation of OsbZIP23 target genes that are thought to provide drought protection^19^. OsOTS1 may mediate the accumulation of similar transcription factors that mediate root growth in rice during salt stress and this might provide a mechanism for salinity tolerance in OsOTS1 overexpressing rice. Thus, our work provides a foundation for further understanding the role of SUMO proteases in growth and development of crop plants such as rice under stress.

## Materials and Methods

### Vector construction

To accommodate the CRISPR/Cas9 system to *Agrobacterium*-mediated plant transformation, we have used the entry vectors and destination vectors from previously described^20^. For generating the CRISPR mutant for OsOTS1, we designed a sgRNA targeting the first exon of *OsOTS1* (Figure 1) and cloned into the entry vector pOs-SgRNA. We designed the sgRNA using the web-based tool CRISPR-P (http://cbi.hzau.edu.cn/crispr/)^21^. The sequences and other data of analysed rice endogenous gene was downloaded from Rice Genome Annotation Project (http://rice.plantbiology.msu.edu/). The OTS1 sequence was analysed for potential sgRNA sequences (20 nucleotides) immediately followed by 5’-NGG (PAM sequence) in the forward and reverse sequence. A pair of DNA oligos were designed and synthesized from Integrative DNA Technology, UK and annealed to generate dimers. These dimers were subsequently ligated upstream of the sgRNA scaffolds in the entry vector within the BsaI restriction site. The binary expression vector pH-Ubi-cas9-7 containing a *Hygromycin* gene and Cas9 driven by the maize ubiquitin promoter was used for genetic transformation.

### Transformation of Rice Seedlings

The vectors were introduced into agrobacterium strain EHA105. For each vector more than 500 embryonic calluses of rice (*Oryza sativa* L. ssp. *japonica cv.* Kitaake) were infected with the agrobacterium according to the previously described protocol with slight modifications^22^. The transgenic plants were selected and regenerated under selection with 50mg/l hygromycin. After two to three weeks of rooting, the T_0_ plants were transplanted into soil and grown in controlled condition in a glasshouse at 28 °C with 10 hrs of light and 25°C with 14 hrs in the dark cycle. The progeny was generated through self-pollination of each individual. We obtained 25 transgenic plants with *OsOTS1* transformants (named as DL-1 to DL-25).

### Genotyping of targeted mutations

Genomic DNA was extracted from the transgenic *O. sativa* plants and wild type plants using the CTAB method. We designed the PCR primers in the flanking regions of the Cas9/sgRNA targets and analysed the targeted mutagenesis by PCR amplification and sanger sequencing. For the regenerated plants, the presence of CRISPR/Cas9 constructs was investigated by PCR with *Cas9* gene primer. For the transformed *O. sativa* plants, the DNA fragments spanning the Cas9/sgRNA target sequences were amplified by PCR and the products were analysed by TA cloning and sequencing. The primers used for PCR amplification are listed in Table S1.

### Chlorophyll contents measurement

Chlorophyll content was measured as previously described^23^. 0.5 g of fresh leaf sample was suspended in 10 ml of 80% Acetone mixed well and kept at room temperature in the dark for 7d. The supernatant was collected after centrifugation at 5000 rpm/min for 15 min. The sample absorbance was recorded at 645 and 663 nm using a spectrophotometer.

### Off target effect detection

The potential off-target sites were predicted using the online tool CRISPR-P (http://cbi.hzau.edu.cn/CRISPR/)^21^ and the top two off-target sites (from OTS2 and OTS3?) for OTS1 were analysed by sequence analysis. The fragment containing the target region (726 bp for OTS2 and 720 bp for OTS3) was first PCR amplified using the primer pair. The sites of each target in five randomly selected mutant lines were examined by site-specific PCR and direct Sanger sequencing. The sequences of the sites and related PCR primers are listed in Supplementary Table S1.

### RNA extraction and Quantitative RT-PCR

Total RNA was isolated using the RNA isolation kit (Sigma). After RNase-free DNase treatment, 2μg RNA was reversibly transcribed using oligo (dT) primer and reverse transcriptase II. Quantitative PCR was performed on the Qiagen Rotor Gene Q, using the Sigma SYBR Green qPCR mix. The results were analysed using software (Qiagen Rotor Gene Q). Each data set had three replicates and the experiments were repeated twice. Primers are listed in supplementary Table S1.

## Acknowledgements

This work was supported by the European Research Council (ERC) Grant and BBSRC. The authors declare no conflict of interest. We would like to acknowledge Dr. Christopher Perin, CIRAD, Montpellier, France for providing the CRISPR/Cas9 entry and destination vectors.

## Author contribution

CZ, AKS and AS conceived and designed research. CZ and AKS conducted experiments. CZ, AKS and AS analysed data. AKS and AS wrote the manuscript. All authors read and approved the manuscript.

---

**Supplementary Table S1** DNA primers used in the study

